# SV40 exploits the Nesprin-2-SUN1-KPNA4 axis for stepwise targeting and entry into the host nucleus to promote infection

**DOI:** 10.64898/2026.03.15.711898

**Authors:** Luke Gohmann, Billy Tsai

**Affiliations:** Department of Cell and Developmental Biology University of Michigan Medical School, Ann Arbor, Michigan, United States; Cellular and Molecular Biology Program University of Michigan Medical School, Ann Arbor, Michigan, United States

## Abstract

Many DNA viruses including polyomaviruses (PyVs) enter the host nucleus to cause infection, although how this is accomplished is unclear. To infect cells, the prototype PyV SV40 targets to the Nesprin-2 outer nuclear membrane protein and enters the nucleus via the nuclear pore complex (NPC). Host factors that function with Nesprin-2 to target SV40 to the nuclear membrane and drive NPC-dependent nuclear entry are unknown. Here we demonstrate that the SUN1 inner nuclear membrane protein acts coordinately with its binding-partner Nesprin-2 to target cytosol-localized SV40 to the nuclear membrane. Strikingly, despite localizing to the perinuclear space, the SUN domain of SUN1 plays a crucial role in Nesprin-2-dependent recruitment of cytosolic SV40. After targeting, SV40 binds to the NPC-associated importin receptor KPNA4, which translocates the virus into the nucleus. Our results reveal how a DNA virus exploits the Nesprin-2-SUN1-KPNA4 axis for stepwise targeting and entry into the nucleus to cause infection.

**Author summary:** Nuclear entry is required for most DNA viruses to cause infection, although the molecular mechanism of this step remains enigmatic. The DNA virus SV40 targets to the nuclear membrane by exploiting the Nesprin-2 outer nuclear membrane protein. In this study, we report that the SUN1 inner nuclear membrane protein functions with Nesprin-2 to target SV40 to the nuclear membrane, followed by nuclear entry of the virus via the action of the KNPA4 importin receptor.

## Introduction

After entry, many DNA viruses enter the host nucleus to cause infection, although how this decisive infection step is accomplished remains unclear. Elucidating the molecular basis of viral nuclear entry may lead to new therapeutic approaches to combat virus-induced diseases and illuminate basic nuclear transport mechanisms. Polyomaviruses (PyV) are DNA viruses that must gain entry into the host nucleus to cause infection (1), although the molecular mechanism of this step is largely enigmatic. Included in the PyV family is the human BK PyV (BKV) responsible for nephropathy and hemorrhagic cystitis (2), JC PyV that triggers a fatal demyelinating disease called progressive multifocal leukoencephalopathy (3), and Merkel Cell PyV that causes an aggressive skin tumor known as Merkel cell carcinoma (4). Because the steps of infection by the prototype PyV—simian virus 40 (SV40)—are similar to human PyVs (5), a coherent understanding of the SV40 infection mechanism will likely provide critical insights into combating PyV-induced human diseases.

Structurally, SV40 is composed of 72 pentamers of the coat protein VP1 that encloses its DNA genome, with each pentamer encasing an internal hydrophobic protein VP2 or VP3 (6, 7). Three forces stabilize the overall viral architecture: the VP1 C-terminus invades a neighboring VP1 pentamer to provide inter-pentamer support, presence of intra-/inter-pentamer disulfide bonds stabilize the viral structure, and calcium ions bound to the virus further strengthen its structure (8). When assembled, SV40 is 50 nm.

During entry, SV40 undergoes receptor-mediated endocytosis to reach the endosome from where it is trafficked to the endoplasmic reticulum (ER) (9–11). In this compartment, ER-resident PDI family members reduce and isomerize the VP1 inter-pentameric disulfide bonds (12, 13) and unfold the invading VP1 C-terminal arms (14), steps that effectively destabilize the viral architecture. These disruptions expose the internal hydrophobic VP2 and VP3 proteins (14, 15), producing a hydrophobic particle that integrates into the ER membrane at specific sites referred to as the ER-foci (15–19). At the ER foci, a cytosolic extraction machinery extracts SV40 into the cytosol (16, 20–23).

Upon escaping to the cytosol, SV40 is disassembled by the bicaudal D2 (BICD2) protein (24), which normally acts as a cargo adaptor for the cellular dynein motor (25). Specifically, BICD2 triggers the partial uncoating of the VP1 pentamers from the intact viral particle to generate a disassembled virus (24). Importantly, this subviral particle is delivered to the nuclear membrane—likely by the dynein-BICD2 motor complex—to initiate entry into the nucleus. The identities of the nuclear membrane components that capture the cytosol-localized SV40 are slowly being unraveled. We previously reported that Nesprin-2, an outer nuclear membrane protein that is a subunit of the linker of nucleoskeleton and cytoskeleton (LINC) nuclear membrane complex, captures cytosol-localized SV40 (26). However, the identities of other host components that act in concert with Nesprin-2 to recruit SV40 to the nuclear envelope, as well as how the virus is subsequently translocated into the nucleus to promote infection, are unknown. In this context, there is evidence that the nuclear localization signal (NLS) of SV40 VP2 and VP3, along with components of the classic nuclear import machinery, supports virus nuclear entry via the nuclear pore complex (NPC) (27, 28).

In this manuscript, we report that SUN1—an inner nuclear membrane protein that is a binding partner of Nesprin-2 (29)-—plays a decisive role in SV40 infection. SUN1 acts coordinately with Nesprin-2 to target cytosol-localized SV40 to the nuclear envelope. Surprisingly, despite localizing at the perinuclear space, the SUN domain of SUN1 promotes Nesprin-2-dependent recruitment of cytosolic SV40 to the nuclear membrane. After targeting, SV40 is transferred to the NPC-associated importin α receptor KPNA4, which subsequently drives the virus into the nucleus to cause infection. Collectively, our data illuminate the stepwise nuclear targeting and entry mechanism of a DNA virus.

## Results

### SUN1 is required for SV40 infection

The LINC complex is composed of Klarischt/ANC-1/SYNE homology (KASH) proteins embedded in the outer nuclear membrane and Sad1/UNC-84 (SUN) proteins embedded in the inner nuclear membrane (29). The KASH and SUN proteins physically interact at the perinuclear space located between the two nuclear membranes, enabling the LINC complex to span the entire nuclear envelop. In mammals, there are six KASH proteins (Nesprin-1-4 and KASH5-6) and five SUN proteins (SUN1-5), in which only SUN1 and SUN2 are ubiquitously expressed (30, 31). Because we previously found that Nesprin-2 recruits cytosol-localized SV40 to the nuclear membrane (26), we hypothesized that SUN1 or SUN2—both established Nesprin-2 binding-partners (29)—also promotes recruitment of SV40 to the nuclear envelope to enable nuclear entry essential for infection.

To test this hypothesis, we first evaluated whether SUN1 or SUN2 is important for SV40 infection. We used siRNA-mediated knockdown (KD) to deplete the nuclear membrane proteins in simian CV-1 cells, which are used classically to study SV40 infection (32). Accordingly, cells are transfected with either the control scrambled (Scr) siRNA, siRNA against SUN1 or SUN2, and the resulting cell extract subjected to SDS-PAGE and Western blotting. We found the level of SUN1 (but not SUN2) decreased markedly using the SUN1 siRNA (Fig. 1A, first and second blots, compare lane 2 to 1); the level of SUN2 modestly increased under SUN1 KD, possibly reflecting a compensatory response. Similarly, the level of SUN2 (but not SUN1) decreased robustly using the SUN2 siRNA (Fig. 1A, first and second blots, compare lane 3 to 1). The control and SUN protein-depleted cells were then infected with SV40 (MOI 1), and at 24 hours post-infection (hpi), stained for appearance of the large T-antigen (T-Ag) in the nucleus by microscopy. T-Ag is a virally-encoded protein that is expressed upon successful arrival of SV40 to the host nucleus and its appearance marks successful infection. Importantly, using this imaging approach, we found SV40 infection is blocked markedly under SUN1 (but not SUN2) KD (Fig. 1B, compare second and third bars to first). By Western blotting (using a T-Ag antibody), we also found that SUN1– (but not SUN2) depletion robustly impaired SV40 infection (Fig. 1C, top blot, compare lane 2 and 3 to 1; the T-Ag band intensity is quantified in 1D), consistent with the microscopy results (Fig. 1B). These findings suggest that SUN1 plays an important role in SV40 infection.

**Figure 1.**
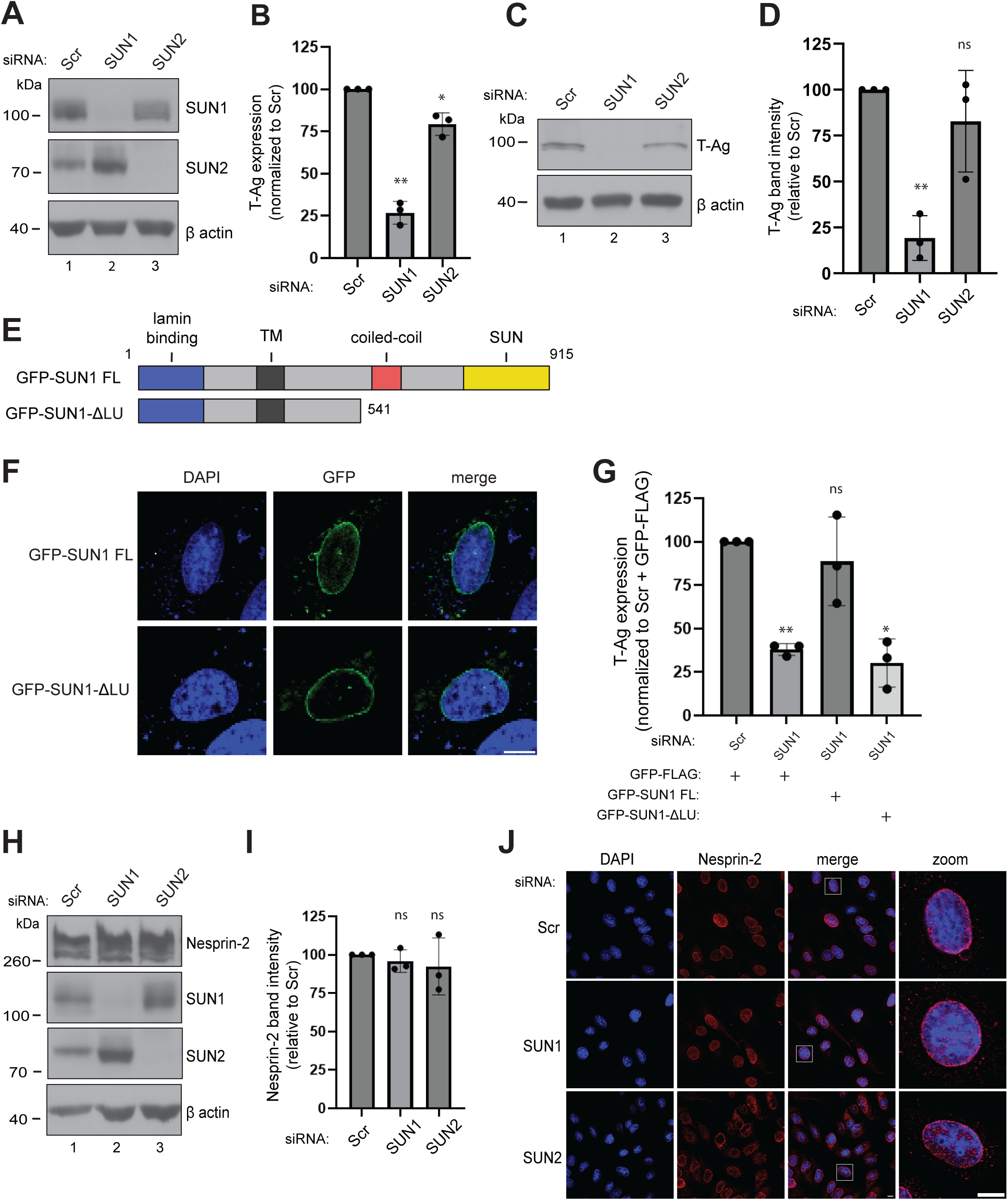
SUN1 is required for SV40 infection. **(A)** CV-1 cells transfected with a scrambled (Scr) control siRNA or siRNA against SUN1 or SUN2 were harvested and the resulting whole cell extracts subjected to SDS-PAGE and immunoblotting with the indicated antibodies. β actin was used as a loading control. **(B)** CV-1 cells transfected with the indicated siRNA were infected with SV40, fixed, and stained for large T antigen (T-Ag). Data were normalized to the Scr control. **(C)** CV-1 cells transfected with the indicated siRNA were infected with SV40 and the resulting whole cell extracts were subjected to SDS-PAGE and immunoblotting. **(D)** The T-Ag band intensity in C was quantified by the FIJI software. Data were normalized to the Scr control. **(E)** Schematic of full-length (FL) SUN1 containing an N-terminal GFP tag (GFP-SUN1 FL), and a truncated SUN1 lacking the coiled-coil and SUN domain with also an N-terminal GFP tag (GFP-SUN1 ΔLU). **(F)** CV-1 cells transfected with the indicated constructs were fixed and stained for GFP (green) and counterstained with DAPI (blue). Scale bar: 10 µm. **(G)** CV-1 cells transfected with the Scr control siRNA or siRNA against SUN1 were also transfected with either the GFP-FLAG control construct or the indicated GFP-SUN1 construct. Cells were infected with SV40, fixed, and stained to assess T-Ag expression, as in 1B. Only GFP-expressing cells were analyzed and data were normalized to the Scr control with GFP-FLAG. **(H)** CV-1 cells were transfected with either Scr or siRNA against SUN1 or SUN2. The resulting whole cell extracts were subjected to SDS-PAGE and immunoblotted with the indicated antibodies. **(I)** The Nesprin-2 band intensity in H was quantified by the FIJI software and data were normalized to the Scr control. **(J)** CV-1 cells transfected with either Scr or siRNA against SUN1 or SUN2 were fixed and stained for Nesprin-2 (red) and counterstained with DAPI (blue). * p ≤ 0.05; ** p ≤ 0.01; ns= not significant. Scale bar: 10 µm

We next performed a KD-rescue experiment to determine if the block in SV40 infection due to the SUN1 siRNA results from SUN1 depletion, as well as to determine whether interaction between SUN1 and Nesprin-2 is required to promote virus infection. SUN1 is a single-pass transmembrane (TM) protein that contains an N-terminal lamin binding domain oriented in the nucleoplasm, a TM domain, followed by a coiled-coil and a C-terminal SUN domain that face the perinuclear space (Fig. 1E). In the rescue experiments, we used a previously published GFP-tagged full-length SUN1 construct (GFP-SUN1 FL;(33)) and a GFP-tagged SUN1 deletion construct lacking the SUN (and coiled-coil) domains (GFP-SUN1 ΔLU;(33)) because the SUN domain is responsible for Nesprin-2 binding (31); both constructs are designed to be resistant to the SUN1 siRNA. As previously reported (33), both GFP-SUN1 FL and GFP-SUN1 ΔLU are localized to the nuclear membrane (Fig. 1F, top and bottom rows). Importantly, we found that expression of GFP-SUN1 FL, but not GFP-SUN1 ΔLU, rescued the block in SV40 infection under SUN1 KD (Fig. 1G, compare third and fourth bars to second bar). These findings demonstrate that the block in SV40 infection due to the SUN1 siRNA is due to loss of SUN1 and not to unintended off-target effects. They also indicate that intact SUN1-Nesprin-2 interaction is essential for SUN1 to promote virus infection.

The Nesprin-2 level was unchanged under SUN1 (or SUN2) KD (Fig. 1H, top blot, compare lane 2 and 3 to 1; the Nesprin-2 band intensity is quantified in 1I), and Nesprin-2 maintained its localization to the nuclear membrane in SUN1-depleted cells (Fig. 1J, compare middle to top row; see zoom). These control experiments demonstrate that the impaired SV40 infection in SUN1-depleted cells is not caused by a reduced Nesprin-2 level or mislocalized Nesprin-2. Collectively, our findings establish SUN1 is required for SV40 infection.

### SUN1 promotes nuclear targeting of SV40

Nesprin-2 forms a complex with SUN1 or SUN2 on the nuclear membrane (31), raising the possibility that the Nesprin-2-SUN1 or Nesprin-2-SUN2 protein complex is responsible for nuclear targeting of SV40. To test whether SUN1 or SUN2 plays a role in targeting SV40 to the nuclear membrane, we used a microscopy assay which was used previously to establish a role of Nesprin-2 in SV40 nuclear targeting (26). In this assay, we tracked presence of the VP2/3+ signal on the nuclear envelop of SV40-infected cells because during nuclear entry a SV40 particle harboring VP2/3 also contains the viral genome and thus represents an infectious viral particle (26). In these experiments, the transcription inhibitor actinomycin D was included at the time of infection to ensure that only the incoming viral particle, but not newly transcribed/translated virus proteins, is tracked.

Using this assay, we found that whereas the VP2/3+ signal is present on the nuclear membrane of control and SUN2 KD cells (infected with SV40 for 20 h), this signal was absent from the nuclear membrane in SUN1-depleted cells (Fig. 2A, compare top and bottom rows to middle row, see merge; the percentage of cells with the VP2/3+ signal on the nuclear membrane is quantified in Fig. 2B). Of note, the VP2/3+ signal was not present in the nucleoplasm because the cells were infected with SV40 for only 20 h when the virus has only reached the nuclear membrane but not yet entered the nucleoplasm (see Fig. 6G for comparison). These findings indicate that SUN1 (but not SUN2) is responsible for nuclear targeting of SV40. Because depletion of Nesprin-2 also blocked nuclear targeting of SV40 (26), our findings suggest that the Nesprin-2-SUN1—but not the Nesprin-2-SUN2—protein complex recruits cytosol-localized SV40 to the nuclear membrane.

**Figure 2.**
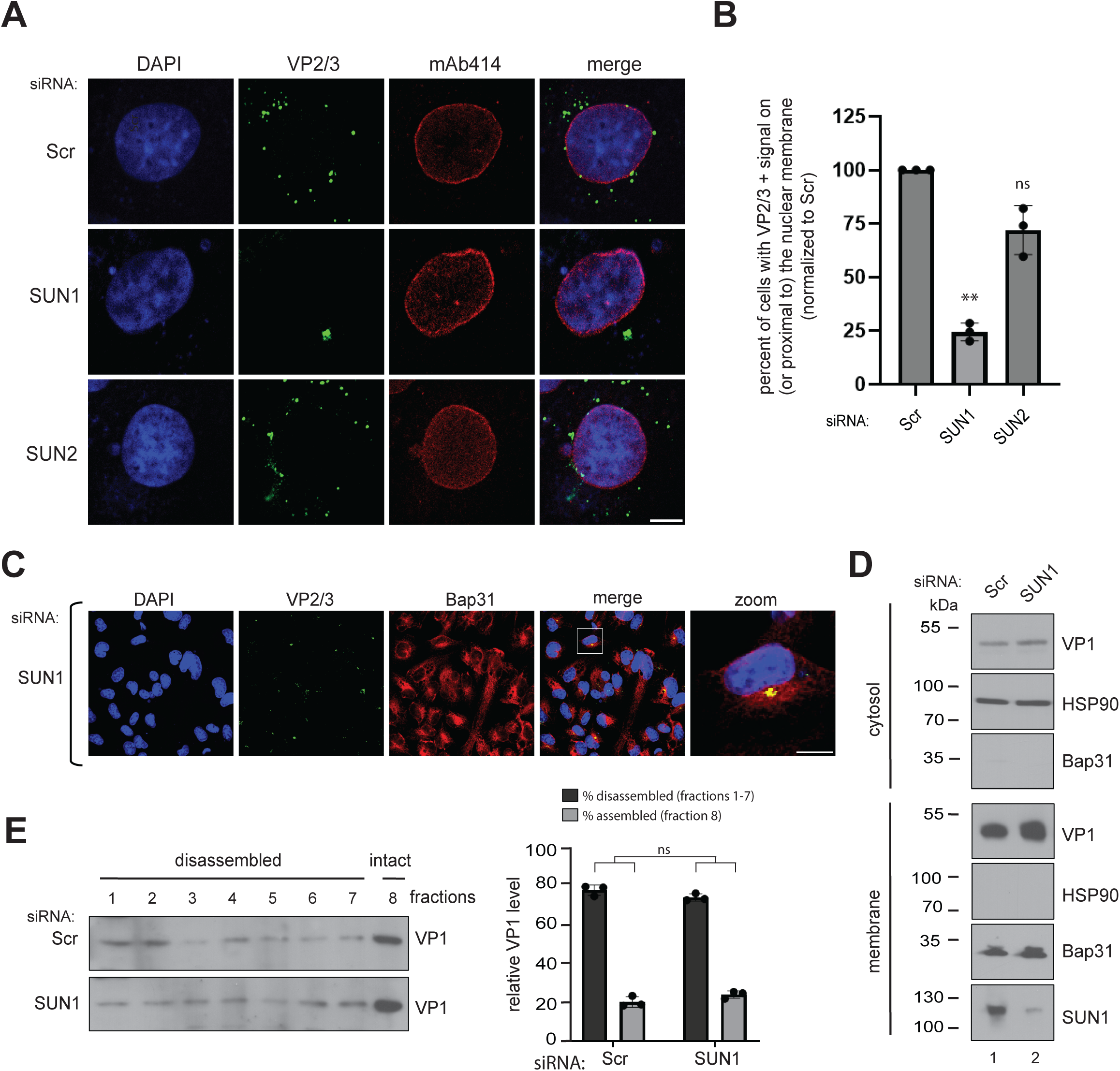
SUN1 promotes nuclear targeting of SV40. **(A)** CV-1 cells transfected with either a scrambled (Scr) control siRNA or siRNA against SUN1 or SUN2 were infected with SV40 in the presence of actinomycin D (ActD). Cells were stained with anti-VP2/3 (green), mAb414 (red), and counterstained with DAPI (blue). Scale bar: 10 µm. **(B)** The percent of cells with a discrete VP2/3+ signal on or proximal to (i.e. within 0.3 µm) the nuclear membrane were quantified and normalized to the Scr control. **(C)** As in A, except stained with Bap31 (red). Scale bar: 10 µm. **(D)** CV-1 cells transfected with the indicated siRNA were infected with SV40 and processed using the ER-to-cytosol transport assay as described in ((20)). The resulting cytosol and membrane fractions were subjected to SDS-PAGE followed by immunoblotting with the indicated antibodies. (**E)** The cytosol fraction from D was layered on a discontinuous sucrose gradient and centrifuged to generate individual fractions (see Materials and Methods). Fractions were subjected to SDS-PAGE followed by immunoblotting for VP1. The VP1 signal from fractions 1-7 (representing disassembled virus) and fraction 8 (representing assembled virus) was quantified and normalized to the Scr control. ** p ≤ 0.01; ns= not significant.

Under SUN1 KD, the VP2/3+ signal is not randomly diffused in the cytosol but instead accumulated in a discrete punctum that colocalized with the ER-foci marker Bap31 (Fig. 2C), similarly observed under Nesprin-2 KD (26). The ER-foci represent sites where SV40 escapes into the cytosol and is disassembled by BicD2 (24). To test if SV40 at the ER-foci (under SUN1 KD) is on the cytosolic side of the ER-foci, we performed an established ER-to-cytosol transport assay designed to monitor arrival of SV40 to the cytosol (20). Briefly, SV40-infected cells were treated with a low concentration of a mild detergent (i.e. 0.01% digitonin) to selectively permeabilize the plasma membrane without compromising the integrity of the internal organelles. Cells are then centrifuged to generate a pellet fraction which contains membranes, and a supernatant fraction that harbors cytosolic proteins, as well as SV40 that reached the cytosol. We found that the level of VP1 in the cytosolic fraction was unaffected under SUN1 KD when compared to the control (Fig. 2D, top blot, compare lane 2 to 1). This finding demonstrates that depletion of SUN1 did not impair SV40 arrival to the cytosol and suggest that SV40 at the ER-foci is located on the cytosolic side of this membrane penetration structure.

Because SV40 in the cytosol is disassembled (24, 34, 35), we used a previously-established sucrose gradient disassembly assay (24, 35, 36) to evaluate whether cytosol-localized SV40 (generated in the ER-to-cytosol transport assay) was disassembled. Using the disassembly assay (see Materials and Methods), we found SV40 in the cytosol fraction from control and SUN1 KD cells was disassembled with equal efficiency (Fig. 2E, compare top to bottom blots; the levels of disassembled and assembled SV40 are quantified in the right graph). These biochemical analyses demonstrate that SV40 reaches the cytosol and is disassembled in SUN1-depleted cells. Hence, the block in nuclear targeting of SV40 observed under SUN1 KD (Fig. 2A and 2B) cannot be attributed to impaired arrival of the virus to the cytosol, further strengthening the idea that SUN1 promotes nuclear targeting of SV40 only after arrival of the virus to the cytosol.

### SUN1 and Nesprin-2 act coordinately to recruit SV40

Because our findings indicate that SUN1 is required to target SV40 to the nuclear membrane (Fig. 2), we hypothesize that SUN1 recruits SV40 to the nuclear membrane by interacting with the virus. We used co-immunoprecipitation (coIP) to determine whether SUN1 associates with SV40. When extracts derived from mock– or SV40-infected cells were incubated with a SUN1 (or a control IgG) antibody and the samples subjected to IP, the SV40 viral genome was found in the SUN1 (but not IgG) precipitated material (Fig. 3A, top gel, compare lane 1 to 3), indicating SUN1 associates with the SV40 genome. Additionally, when extracts derived from virus-infected cells expressing FLAG-tagged SUN1 (SUN1-3xFLAG), SUN2 (SUN2-3xFLAG), or the control GFP (GFP-FLAG) were subjected to FLAG IP, VP1 co-precipitated only with SUN1-3xFLAG but not SUN2-3xFLAG or GFP-FLAG (Fig. 3B, top blot, compare lane 3 to 2 and 1); similar results were observed for the SV40 genome (Fig. 3C). These data demonstrate that SUN1 interacts with SV40 harboring the viral genome, which is required for infection.

**Figure 3.**
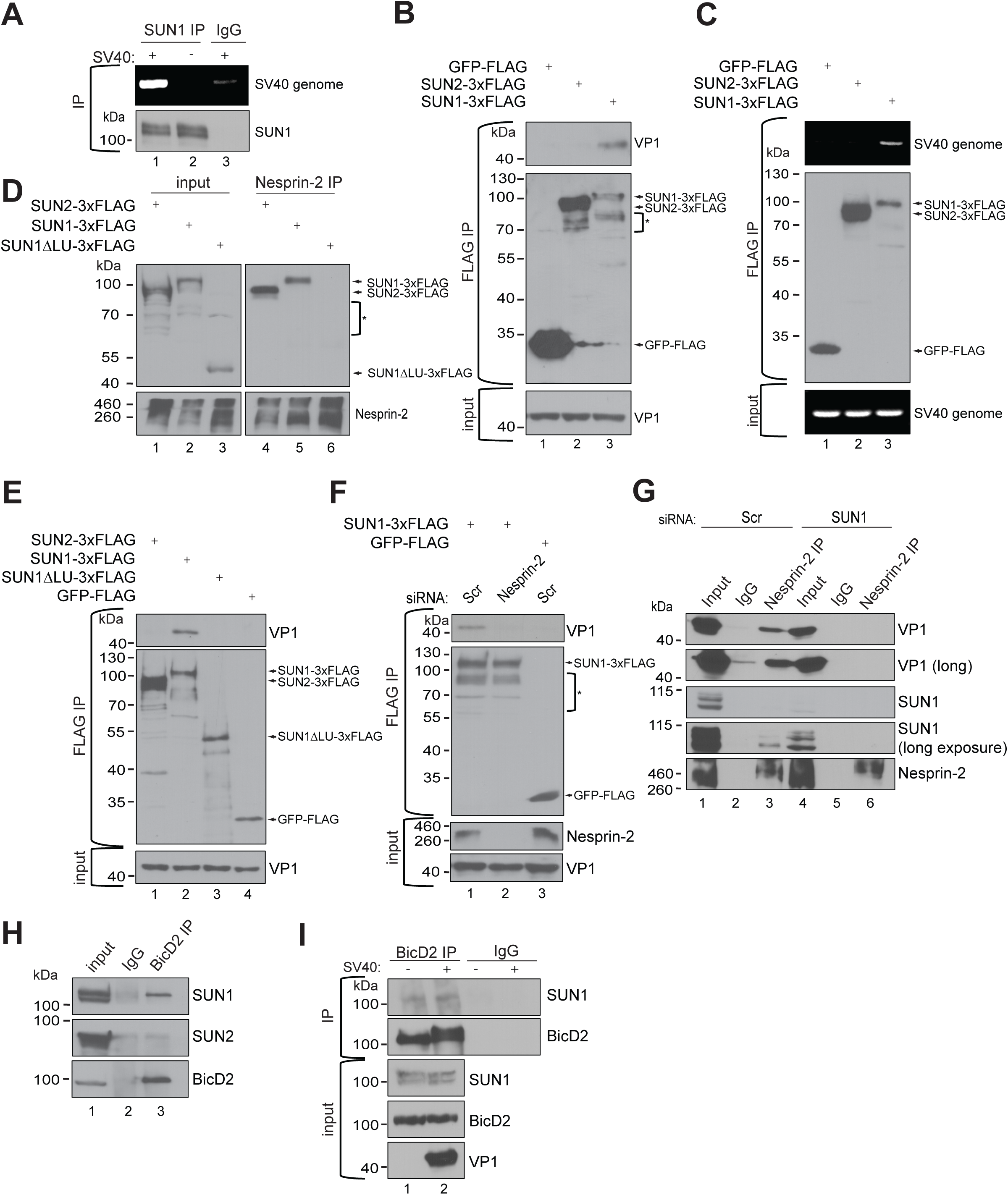
SUN1 and Nesprin-2 act coordinately to recruit SV40. **(A)** Endogenous SUN1 was immunoprecipitated (IPed) from whole cell extracts derived from SV40-infected CV-1 cells. Bound DNA was eluted from the precipitated material and subjected to PCR to detect the SV40 genomic DNA. **(B)** CV-1 cells expressing the indicated FLAG-tagged constructs were infected with SV40. FLAG-tagged proteins were IPed from the resulting whole cell extracts, the precipitated samples eluted and subjected to SDS-PAGE followed by immunoblotting with the indicated antibodies. * indicates degraded SUN proteins. **(C)** As in B, except the precipitated samples were subjected to PCR to detect the SV40 genome. **(D)** Nesprin-2 was IPed from whole cell extracts derived from CV-1 cells expressing the indicated constructs. The precipitated samples were subjected to SDS-PAGE and immunoblotting with the indicated antibodies. * indicates degraded SUN proteins **(E)** As in B, except SUN1 ΔLU-3xFLAG was also used. **(F)** CV-1 cells transfected with the indicated siRNA and construct were infected with SV40. FLAG-tagged proteins were IPed from the resulting whole cell extracts and processed as in B. * indicates degraded SUN1 protein. **(G)** CV-1 cells transfected with the indicated siRNA were infected with SV40. The resulting whole cell extracts were subjected to Nesprin-2 IP and the precipitated samples subjected to SDS-PAGE followed by immunoblotting with the indicated antibodies. **(H)** Endogenous BicD2 was IPed from whole cell extracts derived from CV-1 cells and the precipitated samples subjected to SDS-PAGE and immunoblotting. **(I)** Endogenous BicD2 was IPed from whole cell extracts derived from mock or SV40-infected CV-1 cells. The precipitated samples were subjected to SDS-PAGE and immunoblotting.

Nesprin-2 is located at the outer nuclear membrane with its N-terminus facing the cytosol while SUN1 is located at the inner nuclear membrane with its N-terminus oriented towards the nucleoplasm. The topology of the Nesprin-2-SUN1 protein complex suggests that SUN1 interaction with cytosol-localized SV40 might be indirect and instead depend on Nesprin-2. To test this, we generated a FLAG-tagged SUN1 deletion construct lacking the coiled-coil and SUN domains (SUN1 ΔLU-3xFLAG), which should not bind to Nesprin-2. As expected, IP of Nesprin-2 pulled down SUN1-3xFLAG (and SUN2-3xFLAG) but not SUN1 ΔLU-3xFLAG (Fig. 3D, top blot, compare lanes 5 and 4 to 6). Importantly, by coIP, we found that in contrast to full-length SUN1-3xFLAG, SUN1 ΔLU-3xFLAG cannot pull down SV40 (Fig. 3E, top blot, compare lane 3 to 2); precipitation of GFP-FLAG or SUN2-3xFLAG did not pull down the virus (Fig. 3E, top blot, lanes 4 and 1), as anticipated (Fig. 3B). This finding suggests that SUN1 must properly interact with Nesprin-2 to recruit SV40. To further evaluate whether the SUN1-SV40 interaction depends on Nesprin-2, we depleted Nesprin-2 and found that SUN1-3xFLAG can no longer bind to SV40 (Fig. 3F, top blot, compare lane 2 to 1), further strengthening the idea that SUN1 depends on Nesprin-2 for SV40 binding.

We then performed a reciprocal experiment to assess whether the Nesprin-2-SV40 interaction—observed previously (26)—might rely on SUN1. IP of Nesprin-2 pulled down SV40 in control but not SUN1-depleted cells (Fig. 3G, top two blots, compare lane 3 to 6). This finding indicates that the Nesprin-2-SV40 interaction depends on SUN1, suggesting that SUN1 plays an important role in enabling Nesprin-2 on the nuclear membrane to capture cytosol-localized virus. Together, the bindings experiments suggest that SUN1 and Nesprin-2 act coordinately to recruit cytosol-localized SV40 to the nuclear membrane, preparing the virus for nuclear entry to cause infection.

Why does SV40 exploit the Nesprin-2-SUN1, but not the Nesprin-2-SUN2, protein complex to target to the nuclear membrane? After SV40 reaches the cytosol and is disassembled by the dynein cargo adaptor BicD2 (24), the subviral particle is likely delivered by BicD2, along with the dynein motor, to the nuclear membrane. Because BicD2 interacts to Nesprin-2 (37, 38), we reasoned that BicD2 ought to bind to the Nesprin-2-SUN1 and Nesprin-2-SUN2 protein complex. IP of BicD2 pulled down SUN1 but not SUN2 (Fig. 3H, compare top and middle blots, lane 3) and the SUN1-BicD2 interaction is not affected by SV40 infection (Fig. 3I, top blot, compare lane 2 to 1). These results indicate that BicD2 interacts with only the Nesprin-2-SUN1 but not the Nesprin-2-SUN2 protein complex. Association of BicD2 with the Nesprin-2-SUN1 complex is likely a critical physical feature that enables cytosol-localized SV40 to target to this nuclear membrane complex (see Discussion).

### The SUN domain of SUN1 is crucial to promote SV40 binding and infection

How might the Nesprin-2-SUN1 (but not the Nesprin-2-SUN2) complex support SV40 infection? Notably, a recent report found that the SUN domain of SUN1 promotes Nesprin-2 interaction with microtubules and the microtubule-dependent dynein motor, whereas the SUN domain of SUN2 promotes Nesprin-2 engagement with actin filaments (39–41), which relies on myosin motors (42). Because the BicD2-dyenin motor complex is important for ER-to-nucleus transport of SV40 (24, 25), we suspected that the SUN domain is the defining feature in SUN1 crucial for virus infection. To test this, we used chimeric SUN proteins with swapped SUN domains. Specifically, we generated FLAG-tagged SUN1 with the SUN domain of SUN2 (FLAG-SUN1-S2SUN) and SUN2 with the SUN domain of SUN1 (FLAG-SUN2-S1SUN) (Fig. 4A); we also generated full-length wildtype FLAG-tagged SUN1 and SUN2. Of note, because these SUN proteins are murine proteins, they are resistant to the SUN1 siRNA designed to deplete simian SUN1. As controls, we found by confocal microscopy imaging that the wildtype and chimeric SUN proteins are properly localized to the nuclear membrane (Fig. 4B).

**Figure 4.**
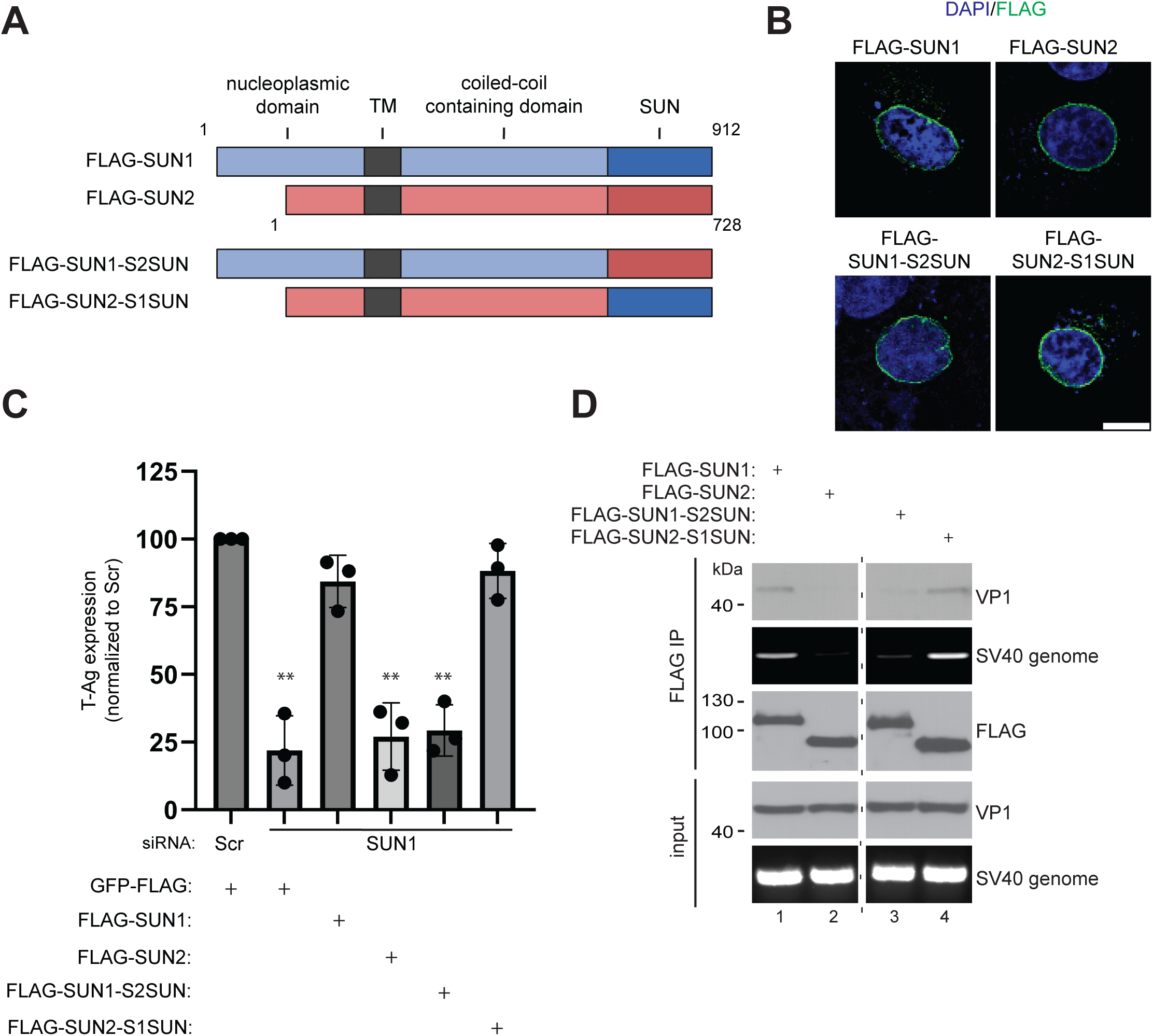
The SUN domain of SUN1 is crucial to promote SV40 binding and infection. **(A)** Schematic of full-length N-terminal FLAG-tagged SUN1 and SUN2, and of chimeric SUN proteins with swapped SUN domains. **(B)** CV-1 cells transfected with the indicated construct were fixed, stained for FLAG (green), and counterstained with DAPI (blue). Scale bar: 10 µm. **(C)** CV-1 cells transfected with the indicated construct and siRNA were infected with SV40, fixed, and stained for T-antigen, FLAG, and counterstained with DAPI. The percentage of transfected cells expressing T-antigen was quantified and normalized to the GFP+Scr control condition. **(D)** CV-1 cells transfected with the indicated construct were infected with SV40. FLAG-tagged proteins were IPed from the resulting whole cell extracts. The precipitated materials were subjected to PCR to detect SV40 genomic DNA or subjected to SDS-PAGE and immunoblotting. The dotted line indicates intervening lanes were removed with adjacent lanes spliced from the same immunoblot. ** p ≤ 0.01

We then used these chimeric (and wildtypes) proteins in KD-rescue experiments, as in Fig. 1G. In cells expressing the control protein GFP-FLAG, depletion of SUN1 blocked SV40 infection (Fig. 4C, compare second to first bar), as expected (Fig. 1G). Under SUN1 KD, SV40 infection is restored when FLAG-SUN1 (but not FLAG-SUN2) is expressed (Fig. 4C, compare third and fourth bars to second bar), as anticipated (Fig. 1G). Strikingly, expression of FLAG-SUN2-S1SUN (but not FLAG-SUN1-S2SUN), restored SV40 infection in SUN1-depleted cells (Fig. 4C, compare sixth and fifth bars to second bar). These findings demonstrate that the SUN domain holds a key structural feature in SUN1 that promotes SV40 infection.

Based on these findings, we postulate that the SUN2-S1SUN chimera, similar to wildtype SUN1, should bind SV40. Indeed, when extracts derived from virus-infected cells expressing these constructs were subjected to FLAG IP, VP1 and SV40 viral genome were pulled down with FLAG-SUN2-S1SUN and FLAG-SUN1, but not FLAG-SUN1-S2SUN or FLAG-SUN2 (Fig. 4D, top two blots, compare lanes 1 and 4 to 2 and 3). These results indicate that the SUN domain is critical for SUN1 to recruit SV40 to the nuclear membrane, consistent with this domain serving a vital role in SUN1-dependent SV40 infection (Fig. 4C).

### KPNA4 is important for SV40 infection

After Nesprin-2-SUN1-dependent nuclear targeting, SV40 translocates into the nucleus to cause infection. How this step is achieved is enigmatic, although there is evidence that NPC-mediated nuclear import is involved (27, 43). Importin α is a key component of the nuclear import machinery, binding to the NLS motif of a cellular cargo to drive nuclear entry (44). Intriguingly, of the seven importin α family members (i.e. KPNA1-7), only KPNA4 was reported to interact with Nesprin-2 (45). We therefore tested whether SV40 exploits KPNA4 for infection; as controls, we evaluated whether KPNA1 and KPNA2 also participate in SV40 infection.

Strikingly, using pooled siRNA to individually deplete each of the three importin α family members (Fig. 5A, bottom, compare lane 2 to 1), we found that depletion of KPNA4 (but not KPNA1 or KPNA2) markedly blocked SV40 infection (Fig. 5A, top, compare lane 4 to 1-3; the T-Ag band intensity is quantified in Fig. 5B). We then used three of the four distinct siRNAs (i.e. KPNA4 #1, #2, or #3 siRNA) in the pooled KPNA4 siRNA separately to deplete KPNA4 (Fig. 5C, second blot, compare lanes 2-4 to 1) and again found that SV40 infection was blocked robustly under these conditions (Fig. 5C, top blot, compare lanes 2-4 to 1; quantified in Fig. 5D). Together, these findings demonstrate that KPNA4 is important for SV40 infection. Confocal imaging revealed that KPNA4 is more strongly localized to the nuclear membrane (as assessed by colocalization with the nuclear membrane marker Nesprin-2) than KPNA1 and KPNA2 (Fig. 5E, compare merge image of third to first and second rows; quantification of the colocalization with Nespin-2 is in Fig. 5F). Preferential localization of KPNA4 to the nuclear membrane suggests that this importin receptor may be strategically positioned at the nuclear membrane to facilitate nuclear entry—presumably after nuclear targeting—to promote virus infection.

**Figure 5.**
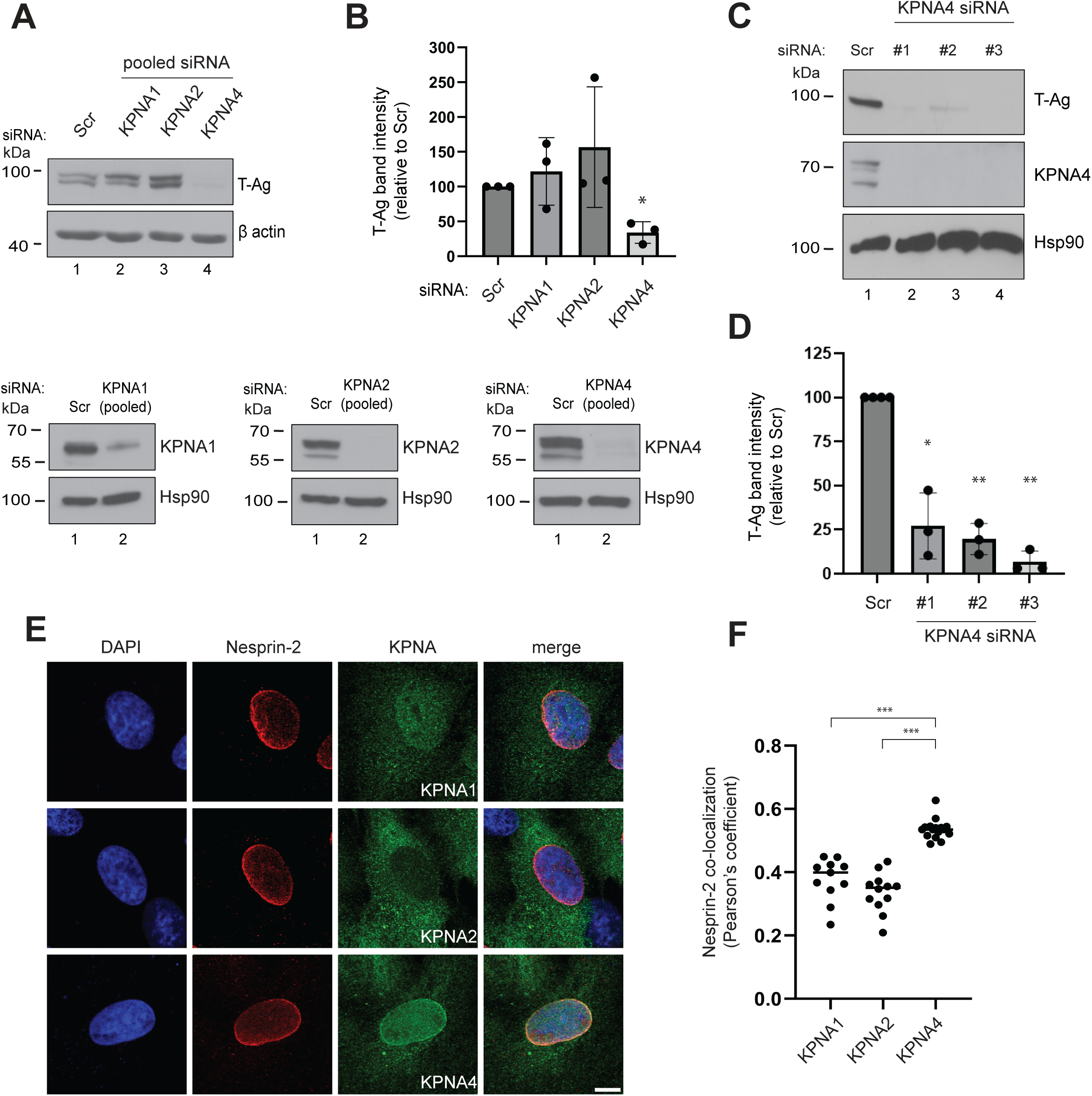
KPNA4 is important for SV40 infection. **(A)** CV-1 cells transfected with the indicated siRNA were infected with SV40 and the resulting whole cell extracts were subjected to SDS-PAGE followed by immunoblotting. The extent of depletion by the KPNA1, KPNA2, and KPNA4 siRNA is shown in the Western blots below. Data were normalized to the Scr control. **(B)** The T-Ag band intensity in A was quantified by the FIJI software. Data were normalized to the Scr control. **(C)** CV-1 cells transfected with the indicated siRNA were infected with SV40 and the resulting whole cell extracts were subjected to SDS-PAGE and immunoblotting. **(D)** The T-Ag band intensity in C was quantified by the FIJI software. Data were normalized to the Scr control. **(E)** CV-1 cells were fixed and stained for Nesprin-2 (red) and either one of the KPNA1, KPNA2, or KPNA4 protein (green) and counterstained with DAPI (blue). Scale bar: 10 µm. **(F)** Pearson’s coefficient was used to quantify colocalization between KPNA1, KPNA2, or KPNA4 with Nesprin-2. Each data point represents one field of view with at least 10 cells. * p ≤ 0.05; ** p ≤ 0.01; *** p ≤ 0.001

### KPNA4 binds to SV40 after Nesprin-2-SUN1-dependent targeting to promote virus nuclear entry

Because our findings indicate that KPNA4 is required for infection, we hypothesize that KPNA4 interacts with SV40 prior to nuclear entry; we used the coIP method to test this idea. We found that IP of KPNA4 using an extract from virus-infected cells pulled down SV40 (Fig. 6A, top blot, lane 3), along with SUN1 and Nesprin-2 (Fig. 6A, second and third blots, lane 3), as expected (45). These results demonstrate that KPNA4 binds to SV40, likely to drive the virus into the nucleus.

We then determined whether the KPNA4-SV40 interaction depends on Nesprin-2-SUN1-mediated SV40 targeting to the nuclear membrane. We found whereas IP of KPNA4 in control cells pulled down SV40 as expected, the KPNA4-SV40 interaction was abolished in SUN1-depleted cells (Fig. 6B, top blot, compare lane 2 to 1). Thus, the KPNA4-SV40 interaction depends on SUN1, indicating that this interaction occurs downstream of Nesprin-2-SUN1. By contrast, the SUN1-SV40 interaction was enhanced under KPNA4 KD (Fig. 6C, top blot, compare lane 2 to 1; quantified in Fig. 6D), suggesting that the virus accumulates on the Nesprin-2-SUN1 complex in KPNA4-depleted cells. Consistent with this, in cells infected with SV40 for 20 h, we found modestly increased colocalization between SV40 (i.e. VP2/3) and SUN1-3xFLAG under KPNA4 KD (Fig. 6E; quantified in Fig. 6F). These data further support the notion that the SV40-KPNA4 interaction occurs after the virus is targeted to the Nesprin-2-SUN1 complex.

**Figure 6.**
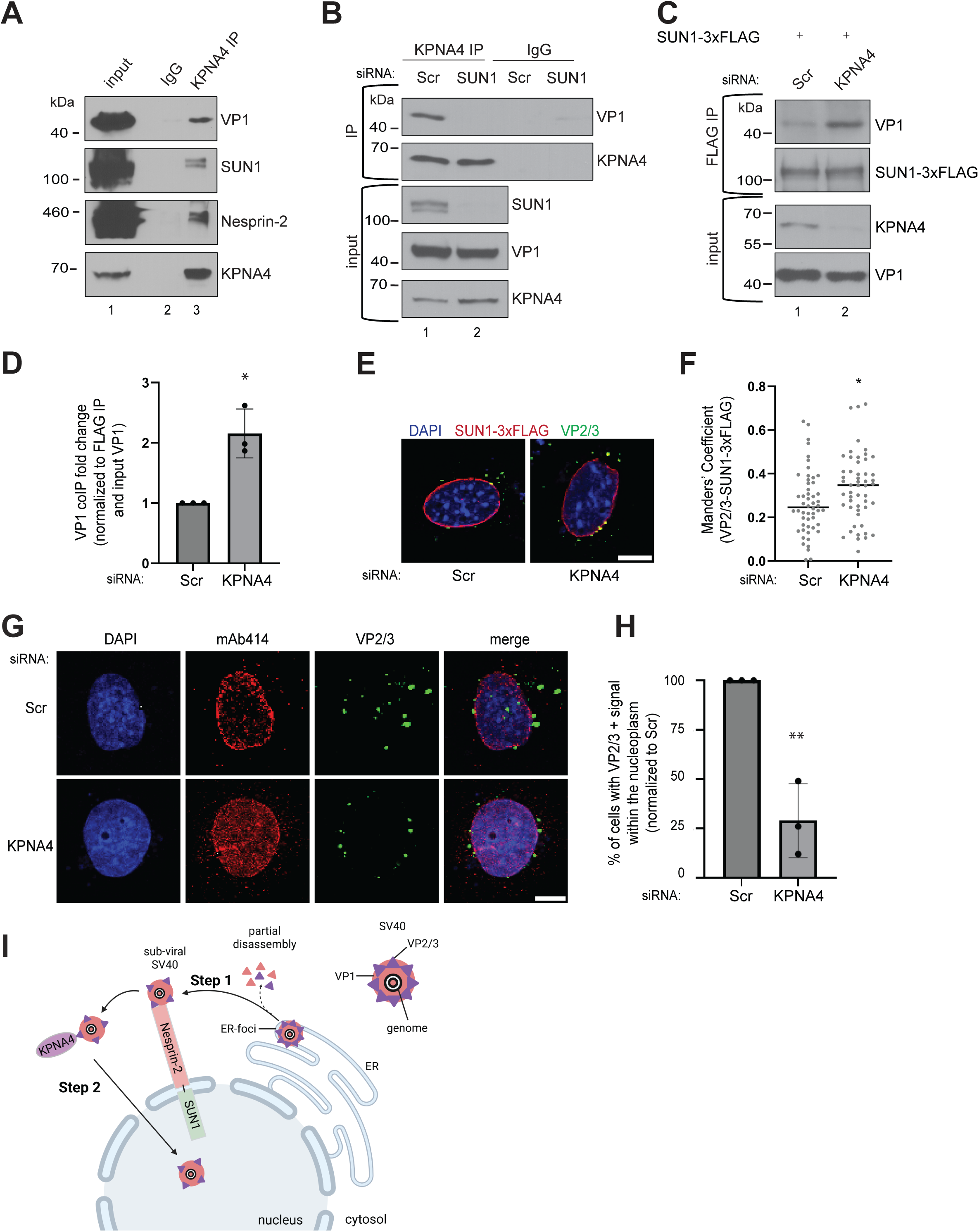
KPNA4 binds to SV40 after Nesprin-2-SUN1-dependent targeting to promote virus nuclear entry. **(A)** Endogenous KPNA4 was IPed from whole cell extracts derived from SV40-infected CV-1 cells. The precipitated samples were subjected to SDS-PAGE and immunoblotting. **(B)** As in A, except CV-1 cells were subjected to SUN1 KD. **(C)** CV-1 cells transfected with the indicated siRNA and SUN1-3xFLAG were infected with SV40. SUN1-3xFLAG was IPed from the resulting whole cell extracts, and the precipitated samples subjected to SDS-PAGE and immunoblotting. **(D)** The VP1 band intensity in C was quantified by the FIJI software. Data were normalized to the Scr condition. **(E)** As in Fig. 2A, except cells were depleted of KPNA4 and stained for FLAG (red), VP2/3 (green), and counterstained with DAPI (blue). Scale bar: 10 µm. **(F)** Manders’ coefficient was used to quantify overlap of VP2/3+ signal on SUN1-3xFLAG. Each data point represents one cell. **(G)** CV-1 cells transfected with the indicated siRNA were infected with SV40. Cells were fixed and stained for mAb414 (red), VP2/3 (green), and counterstained with DAPI (blue). Scale bar: 10 µm. **(H)** The percent of cells with a discrete VP2/3+ signal in the nucleus in G was quantified and normalized to the Scr control condition. **(I)** Model: After SV40 reaches the cytosol from the ER (foci), the viral particle is partially disassembled and targeted to the Nesprin-2-SUN1 nuclear membrane protein complex (Step 1). The virus is then released to the KPNA4 importin α receptor, which translocates SV40 into the nucleus to promote infection (Step 2). Schematic created in biorender.com. * p ≤ 0.05, ** p ≤ 0.01

If KPNA4 acts after targeting of SV40 to the Nesprin-2-SUN1 complex, we envision that KPNA4 promotes entry of the virus into the nucleus. In this scenario, the block in SV40 infection under KPNA4 KD (Fig. 5) is due to impaired nuclear entry. Confocal microscopy revealed that in SV40-infected cells, the level of the VP2/3+ signal in the nucleoplasm decreased significantly under KPNA4 KD (Fig. 6G, compare merge in bottom to top row; the percentage of cells with VP2/3+ signal in the nucleoplasm is quantified in Fig. 6H). In these experiments, cells were infected with SV40 for 36 h so that the virus can reach the nucleoplasm and allow the VP2/3+ signal to be observed inside the nucleus (compare to Fig. 2A). Of note, SV40 can reach the nuclear membrane under KPNA4 KD (Fig. 6H, see zoom in bottom row), in contrast to the fate of SV40 under SUN1 KD (Fig. 2A). Together, these findings support the hypothesis that KPNA4 is required for nuclear entry of SV40.

## Discussion

Although infection for most DNA viruses requires the viral particle to reach the nucleus, how this is accomplished in many instances remains enigmatic. Upon reaching the cytosol from the ER, the DNA virus SV40 is targeted to the nuclear membrane via engaging the Nesprin-2 outer nuclear membrane protein (26). Host factors that assist Nesprin-2 in targeting SV40 to the nuclear membrane and translocation into the nucleus are unknown. In this manuscript, we identify SUN1 as an inner nuclear membrane protein that acts coordinately with Nesprin-2 to target SV40 to the nuclear membrane (Fig. 6I, step 1). The nuclear membrane-associated importin receptor KNPA4 then binds and translocates SV40 into the nucleus to enable infection (Fig. 6I, step 2).

Because the inner nuclear membrane proteins SUN1 and SUN2 are well-established binding partners of Nesprin-2 (29), we evaluated their roles in SV40 infection by using an siRNA-mediated KD approach and found that SUN1 (but not SUN2) promotes virus infection. A rescue strategy revealed that full length SUN1, and not truncated SUN1 which cannot bind to Nesprin-2, restored SV40 infection under SUN1 KD. These findings demonstrate that SUN1 must engage Nesprin-2 to promote SV40 infection, suggesting that an intact Nesprin-2-SUN1 protein complex is important in mediating virus infection.

We then used a previously established nuclear targeting assay and discovered that SUN1 is necessary for targeting SV40 to the nuclear membrane, consistent with a role of Nesprin-2 in this same step (26). Intriguingly, in SUN1-depleted cells, SV40 is not randomly distributed in the cytosol but instead accumulates on the cytosolic side of the ER foci; SV40 is similarly trapped on the cytosolic side of the ER foci under Nesprin-2 KD (26). These results imply that even though the Nesprin-2-SUN1 complex is localized to the nuclear membrane, this protein complex nonetheless impacts trafficking of SV40 from the cytosolic side of the ER foci to the nucleus.

How might a nuclear membrane protein complex control trafficking of SV40 (localized at the cytosolic side of the ER foci) to the nuclear membrane? The Nesprin-2-SUN1 (but not Nesprin-2-SUN2) complex is known to support microtubule-dependent dynein transport (39, 41). In fact, the dynein cargo adaptor BicD2 was shown to bind to Nesprin-2 (37). Because SV40 that reaches the cytosol (from the ER foci) is partially disassembled by BicD2 (24), the BicD2-dynein motor complex likely delivers this viral particle to Nesprin-2-SUN1 on the nuclear membrane. Thus, we postulate presence of a microtubule-dependent pathway that links SV40—localized at the cytosolic side of the ER foci—to the nuclear membrane, and this transport pathway is stabilized by Nesprin-2-SUN1. In this scenario, without either Nesprin-2 or SUN1, the microtubule-dependent transport pathway to the nuclear envelope destabilizes, causing SV40 to accumulate in the cytosolic side of the ER foci. Additional experiments are needed to address this possibility and to investigate how SV40 is retained near the ER membrane instead of undergoing non-productive trafficking to other sites.

Biochemical analyses revealed that SUN1 interacts with infectious SV40 (i.e. with genome) in a Nesprin-2-dependent manner. This suggests that the SV40-SUN1 interaction is indirect and instead mediated by Nesprin-2, consistent with the ability of Nesprin-2 to bind directly to SV40 (26). Surprisingly, we found that the SV40-Nesprin-2 interaction relies on SUN1, even though SUN1 is localized to the inner nuclear membrane and therefore not exposed to the cytosol where the SV40-Nesprin-2 interaction occurs. Thus, SUN1 may regulate the structural conformation of Nesprin-2 to enable efficient cargo (e.g. SV40) recruitment (see below).

A remaining conundrum is why SV40 targets to the Nesprin-2-SUN1 but not Nesprin-2-SUN2 complex. A recent study demonstrated that the SUN domain of SUN1 promotes Nesprin-2 engagement with microtubules and the microtubule-associated dynein motor, while the SUN domain of SUN2 supports Nesprin-2 recruitment of actin filaments that engage myosin motors (39). These observations agree with our data that the dynein cargo adaptor BicD2 selectively binds to the Nesprin-2-SUN1 (but not Nesprin-2-SUN2) complex. By using a KD-rescue strategy, we found that expression of a SUN2 protein harboring the SUN1 SUN domain rescued SV40 infection (in SUN1-depleted cells), whereas expression of a SUN1 protein containing the SUN2 SUN domain did not.

These results align with the biochemical pulldown experiments, which showed that a SUN2 protein harboring the SUN1 SUN domain (but not a SUN1 protein containing the SUN2 SUN domain) can recruit SV40. How the SUN domains of SUN1 and SUN2 promote different actions of Nesprin-2 is surprising given their high sequence similarity (65% identity (46)). However, computational analyses suggest that the SUN domain of SUN1 laterally associates to form high-order oligomers (31), which may cause Nesprin-2 to similarly oligomerize. This may represent a critical feature of the Nesprin-2-SUN1 complex that is used to support microtubule stability essential for dynein-mediated cargo transport. Again, further investigation is required to elucidate how the SUN domains of SUN1 and SUN2 impact the functions of Nesprin-2.

After Nesprin-2-SUN1-dependent targeting to the nuclear membrane, SV40 translocates into the nucleus to initiate infection. We and others have shown that SV40 exploits the NPC for nuclear entry (26, 47, 48). Classic NPC-dependent nuclear entry is mediated by importin α and β family members, which typically form heterodimers (44). During nuclear entry, importin α binds to the NLS of a cargo, while importin β engages different NPC subunits to facilitate passage of the cargo across the NPC channel (44). The importin β1-specific inhibitor importazole potently blocked SV40 infection (26), suggesting that importin β1 plays a critical role in virus infection. However, the identities of the importin α family members that participate in SV40 infection is less clear.

Strikingly, within the seven-member importin α family (i.e. KPNA1-7), only KPNA4 has been reported to bind to Nesprin-2 (45). Indeed, in KPNA4-depleted cells, SV40 infection was markedly impaired, indicating that KPNA4 is important in supporting infection. Furthermore, binding experiments showed that the virus interacts with KPNA4. These results identify an importin α family member that engages and promotes SV40 infection.

To determine whether KPNA4 acts before or after the Nesprin-2-SUN1 complex during nuclear entry of SV40, we found that SUN1-depletion blocked the SV40-KPNA4 interaction, while KPNA4-depletion enhanced the SV40-SUN1 binding. These findings demonstrate that KPNA4 acts after Nesprin-2-SUN1, suggesting that KPNA4 translocates the virus into the nucleus. Consistent with this, confocal imaging revealed that SV40 arrival to the nucleus is robustly impaired under KNPA4 KD without disrupting virus targeting to the nuclear membrane. Together, these data depict a stepwise model of SV40 nuclear entry: cytosol-localized SV40 is first targeted to the nuclear membrane by engaging the Nesprin-2-SUN1 complex, followed by KPNA4-dependent translocation of the virus into the nucleus.

Beyond SV40, the SUN proteins have also been reported to play crucial roles for other viruses. For instance, both SUN1 and SUN2 regulate nuclear entry of HIV (49–51), while SUN2 promotes replication of flaviviruses (52) as well as propagation of HSV-1 (53). Thus, identifying approaches to selectively cripple the functions of these nuclear membrane proteins may offer a potential pan-antiviral therapeutic strategy.

## Materials and Methods

### Cell lines and reagents

CV-1 cells were obtained from ATCC. Cells were grown in complete Dulbecco’s modified Eagle’s medium (DMEM) containing 10% fetal bovine serum, 10 U/mL penicillin, and 10 μg/mL streptomycin (Gibco, Grand Island, NY). DMEM, Opti-MEM and 0.25% trypsin-EDTA were purchased from Invitrogen (Carlsbad, CA). Bovine serum albumin (BSA) and actinomycin D were purchased from Millipore Sigma (St Louis, MO). Phenylmethylsulfonyl fluoride (PMSF) was purchased from Thermo Scientific (Waltham, MA).

### Preparation of SV40

SV40 was prepared using the OptiPrep gradient system (Sigma), as previously described (20). In brief, cells infected with SV40 were lysed in HN buffer (50 mM HEPES pH 7.5, 150 mM NaCl) with 0.5% Brij58 for 30 min on ice. Following centrifugation, the supernatant was layered on top of a discontinuous 20% and 40% OptiPrep gradient. Tubes were centrifuged at 49,500 rpm for 2 h at 4°C in a SW55Ti rotor (Beckman Coulter, Indianapolis, IN). Purified virus was collected from the white interface that forms between the OptiPrep layers and the aliquots stored at –80°C for future use.

### Antibodies

Antibodies, along with the companies that were purchase from and the corresponding catalog numbers, are indicated: SUN1 (Invitrogen, MA5-47231), SUN2 (Proteintech, 27556-1-AP), β actin (Cell Signaling, 4967S), SV40 large T-antigen (Western blot: Santa Cruz Biotechnology, SC-53448; Immunofluorescence: Santa Cruz Biotechnology, SC-147), Nesprin-2 (IP and Immunofluorescence: Invitrogen, MA5-18075; Western blot: Bethyl, A305-393A), SV40 VP1 (Abcam, ab53977), SV40 VP2/3 (Abcam, ab53983), Bap31 (Invitrogen, MA 3002), Hsp90 (Santa Cruz Biotechnology, sc-13119), M2 FLAG (Millipore, F3165-1MG), FLAG (Millipore, F7425), BicD2 (Abcam, ab117818), KPNA1 (Proteintech, 18137-1-AP), KPNA2 (Invitrogen, 108191AP150UL), KPNA4 (Invitrogen, PA518239), Mab414 (Abcam, ab24609).

### Plasmids

GFP-SUN1 FL and GFP-SUN1 ΔLU were gifts from Dr. Derek Walsh (Northwestern University School of Medicine, (33)), SUN1-3xFLAG was a gift from Dr. Nandakumar, (University of Michigan), SUN2-3xFLAG cDNA was a gift from Dr. Grosse (University of Freiburg), and GFP-FLAG was from Chen et al., 2021 ((54)). cDNA for murine SUN1, SUN2, SUN1-S2FLAG, and SUN2-S1FLAG were gifts Dr. Wakam Chang (University of Macau, (39))

### siRNA transfection

All Star Negative (Qiagen, Valencia, CA) was used as a scrambled control siRNA. The following siRNAs were used in this study:

SUN1 (CAGGACGTGTTTAAACCCACGACTT),

SUN2 (GCAGACAUUCCACCCUGCUUUGGUU),

Nesprin-2 (GAGCAUCACUACAAGCAAAUG),

KPNA1 (Dharmacon M-011305-01),

KPNA2 (Dharmacon M-004702-02),

KPNA4 (Dharmacon M-017477-00),

KPNA4 #1 (GAACAAAUACAGAUGGUAA),

KPNA4 #2 (GAAGACAUCUACAAAUUGG),

KPNA4 #3 (GAAGAUAUCUGUGAAGACU)

### DNA transfection

For CV-1 cells, plasmids were transfected into 50% confluent cells using the FuGENE HD (Promega, Madison, WI) transfection reagent. DNA was allowed to express for at least 24 h prior to experimentation.

### Immunofluorescence and confocal microscopy

Cells were grown on No.1 glass coverslips and fixed with 4% formaldehyde for 15 min. Cells were then permeabilized in PBS with 0.2% Triton X-100 for 5 min and blocked with 5% milk containing 0.02% Tween-20. Primary antibodies were incubated in milk for 1 h at room temperature. Coverslips were then washed 6x in milk and incubated with Alexa Fluor secondary antibodies (Invitrogen) for 30 min at room temperature. Coverslips were washed 3x in milk and 3x in PBS, then mounted using mounting media with DAPI (Abcam). Images were taken on either a Leica Stellaris 5 confocal microscope or a Zeiss LSM 780 confocal laser scanning microscope. FIJI software was used for image processing and analysis. At least 100 cells were imaged per condition for each experiment. Z-stack images were taken for co-localization experiments. At least three independent replicates were quantified for each experiment and a standard Student’s t test was used to determine statistical significance. Scale bars are 10 μm.

### Immunoprecipitation

Cells were lysed in RIPA buffer (50 mM Tris pH 7.4, 150 mM NaCl, 0.25% sodium deoxycholate, 1% NP40, 1 mM EDTA) with 1 mM PMSF and 10 mM N-ethylmaleimide (NEM) (Millipore) for 15 min on ice. Cells were then centrifuged at 13,000 rpm for 10 min at 4°C. Input was collected from the resulting supernatant before 2 μg of the indicated antibody was added to the lysate and rotated overnight at 4°C. The next day, either Protein G or Protein A Dynabeads (Thermo Scientific) or Magnetic Beads (New England Biolabs) were washed 3x with RIPA buffer and rotated with the lysate at 4°C for 1 h, respectively. The tubes were then collected on a magnet and the beads washed 3x with RIPA buffer. For genome extraction, 40 μL of RIPA with 1% SDS was added to the beads and vortexed at room temperature for 5 min. Of this, 20% (8 μL) was removed, 40 μL of PB Buffer (Qiagen) added and the samples run through a Qiagen column. DNA was extracted according to the miniprep protocol. SDS sample buffer was added to the remaining IP lysate and the beads boiled for 10 min at 95°C. Samples were then run on an SDS-PAGE gel and protein levels determined by immunoblotting. For FLAG IPs, anti-FLAG M2 conjugated magnetic beads (Sigma) were added to the lysate and rotated at 4°C for 30 minutes. Beads were then washed 2x with RIPA buffer, 1x with RIPA buffer with 300 mM NaCl, and 1x with RIPA buffer with 500 mM NaCl. Beads were then eluted with FLAG peptide (Pierce). Samples were then processed as described above.

### ER-to-cytosol transport assay

This assay follows a previous protocol ((20)), with minor modifications. Briefly, CV-1 cells were infected with SV40 (MOI 5). 12 hpi, cells were semi-permeabilized with a HNP buffer (50 mM Hepes pH 7.5, 150 mM NaCl and 1 mM PMSF) containing 0.05% digitonin at 4° C for 10 min. The sample was centrifuged at 16,100x g for 10 min at 4° C, and the supernatant (cytosol) and pellet (membrane) fractions were collected.

### SV40 disassembly assay

Cytosolic fraction (generated from the ER-to-cytosol transport assay) were layered over a 20-30-40% discontinuous sucrose gradient. Samples were centrifuged for 30 min at 4°C at 50,000 rpm. From the top, 25 μL aliquots were collected and subjected to SDS-PAGE followed by immunoblotting.

### PCR of viral genome

SV40 genomic DNA was amplified using the KOD Hot Start DNA polymerase (MilliporeSigma) with primers against the viral genome (Fwd, 5’ – GCAGTAGCAATCAACCCACA-3’; Rev, 5’ –CTGACTTTGGAGGCTTCTGG-3’). Amplified DNA was subsequently run on an agarose gel to detect viral DNA.

### Quantification of Western blots

All Western blots were developed on film and quantified using the FIJI software. For disassembly experiments, the signal from disassembled virus (fractions 1–7) was compared to that of intact virus (fraction 8) normalized to the total signal (fractions 1–8). For T-antigen experiments, the T-antigen signal was normalized against the loading control (β actin or Hsp90) and against the Scr control condition. For IP experiments, the IP signal was normalized to the input signal and to the total immunoprecipitated material. At least three independent replicates were quantified for each experiment and a standard Student’s t test was used to determine statistical significance.

## Statistical analysis

All data obtained from at least three independent experiments (biological replicates) were combined for statistical analyses. Results were analyzed using Student’s two-tailed *t* test. Data are represented as the mean values, and the error bars represent means ± SD from at least three biological replicates. The significance was determined by p-value, where * p ≤ 0.05; ** p ≤ 0.01; *** p ≤ 0.001, and ns= not significant.

## Acknowledgements

We would like to thank members of the Tsai lab, especially Drs. Tai-Ting Woo and Yu-Jie Chen, for their insights during the course of this project. BT is supported by NIH R01 AI179566. LG is supported by NIH 1T32GM145470.

